# Altered Stool Microbiota of Infants with Cystic Fibrosis Shows Reduction in Genera Associated with Immune Programming

**DOI:** 10.1101/342782

**Authors:** Katherine M. Antosca, Diana A. Chernikova, Kathryn L. Ruoff, Kewei Li, Margaret F. Guill, Todd A. MacKenzie, Dana B. Dorman, Lisa A. Moulton, Molly A. Williams, Brian J. Aldrich, Irene H. Yuan, Margaret R. Karagas, George A. O’Toole, Juliette C. Madan

## Abstract

Previous work from our group indicated a connection between the gastrointestinal microbiota of infants and children with cystic fibrosis (CF) and airway disease in this population. Here we examine the stool microbiota of infants with CF and from the general population who did not have CF over the first year of life. CF children had reduced gastrointestinal *Bacteroides* and *Bifidobacterium* beginning in infancy, even after adjusting for antibiotic treatment. We also identify several metabolic pathways that are enriched or under represented among the microbial communities in the stool of these young patients with CF as compared to children without CF. In vitro studies demonstrated that exposure of the apical face of a polarized Intestinal cell line to *Bacteroides thetaiotaomicron* significantly reduced production of IL-8 secreted from both the apical and basolateral face of these cells, suggesting a mechanism whereby changes in the intestinal microflora could impact systemic inflammation. This work further establishes a link between gastrointestinal microbiota, systemic inflammation and airway disease, and presents the opportunity for therapeutic probiotic interventions.

**Significance statement:** There is a surprising link between gastrointestinal microbial communities and airway disease progression in CF. Here we show that infants with CF ≤1 year of age show a distinct stool microbiota compared with children of a comparable age from a general population cohort. We detect associations between stool microbes and airway exacerbation events in the cohort of infants with CF, and in vitro studies provide a possible mechanism for this observation. These data argue that current therapeutics do not establish a healthy-like gastrointestinal microbiota in young patients with CF, and we suggest that interventions that direct the gastrointestinal microbiota closer to a healthy state may provide benefit to these patients.

## Text

Cystic fibrosis (CF) is an autosomal recessive disease that is known to significantly alter the microenvironments in the airways, intestine and other organs, and is associated with systemic complications such as chronic infections, pancreatic insufficiency, and ultimately premature morbidity and mortality (1-6). While a great deal of attention has been focused on therapies to treat the chronic airway infections that are often the proximal cause of mortality for these patients, it is the gastrointestinal complications that are at the forefront for CF patients in early life (7-10). Young patients with CF experience difficulty in absorbing nutrients secondary to meconium ileus, pancreatic insufficiency and small bowel bacterial overgrowth (10-13). Poor weight gain is common, resulting in the need for a high calorie diet starting early in life and treatment with pancreatic enzyme replacement therapy (14).

Recent work has suggested an unexpected connection between the gastrointestinal tract and airway function in young patients with CF. Our group has shown that select genera (e.g., *Enterococcus*, *Escherichia*) colonizing the airway of CF patients tend to first appear in the intestines (15). Furthermore, breastfed infants showed a longer period of pulmonary health and stability prior to their first pulmonary exacerbation, with 20% of breastfed children having no exacerbation over the first 30 months of life, compared to exclusively formula-fed infants, all of whom suffered exacerbations by 9 months (16). Given the impact of breastfeeding on intestinal microbiota composition (17-19), one possible explanation for these findings in CF is an effect of intestinal microbial communities on airway disease outcomes. Consistent with this idea, an analysis of a small group of patients (n =13, ages 0-34 months) showed that community composition in the intestine, rather than the airway microbiota, was significantly associated with pulmonary exacerbations prior to age 6 months (16).

Here we further probe the relationship between the intestinal microbiota utilizing the Dartmouth Cystic Fibrosis Infant and Children’s Cohort compared with a subset of infants from a general population cohort -- the New Hampshire Birth Cohort Study. Using these two cohorts, we characterize the differences between the intestinal microbiota of patients with CF and the general population over the first year of life, both to understand the differences in the microbiota associated with the intestinal tract in these two populations beginning in infancy, as well as to determine if there is indeed a mechanistic link between intestinal microbiota and airway disease outcomes through cell culture studies.

## Results

### Patient population

In this study, we analyzed 14 patients with CF and 20 non-CF infants at 4 months and 13 CF patients and 45 non-CF infants at 1 year. Of the subjects with CF, 8 had paired samples analyzed at 4 months and 1 yr, and 6 samples were unique at the 2 time points. The non-CF cohort were randomly selected from a larger cohort of children recruited through the New Hampshire Birth Cohort Study for whom stool samples were collected at approximately 4 months and one year of age (20-22). There was no overlap with the non-CF subjects analyzed at 4 months and 1 year. DNA was extracted from stool samples and analyzed for microbial communities as described in the Materials and Methods.

### The alpha diversity of stool communities is altered in infants with CF in the first year of life

To examine the impact of CF on community (alpha) diversity we calculated the Shannon Diversity Index for infants with CF and without CF at 4 months and one year of age (Figure 1). Two key differences could be identified. First, at 4 months, the alpha diversity of the microbial community in the stool of infants with CF was significantly higher than that of those subjects without CF (p = <0.001). No observable change in diversity occurred between 4 months and 1 year in CF infants (p = 0.87); in contrast, the α-diversity did increase during this period in non-CF infants (p < 0.0001). Interestingly, the α-diversity of CF and non-CF children did not significantly differ from each other at 1 year. These data indicate a significant difference in the microbial diversity associated with the stool of infants with CF versus the comparison group of non-CF children as early as 4 months of age.

**Figure 1.**
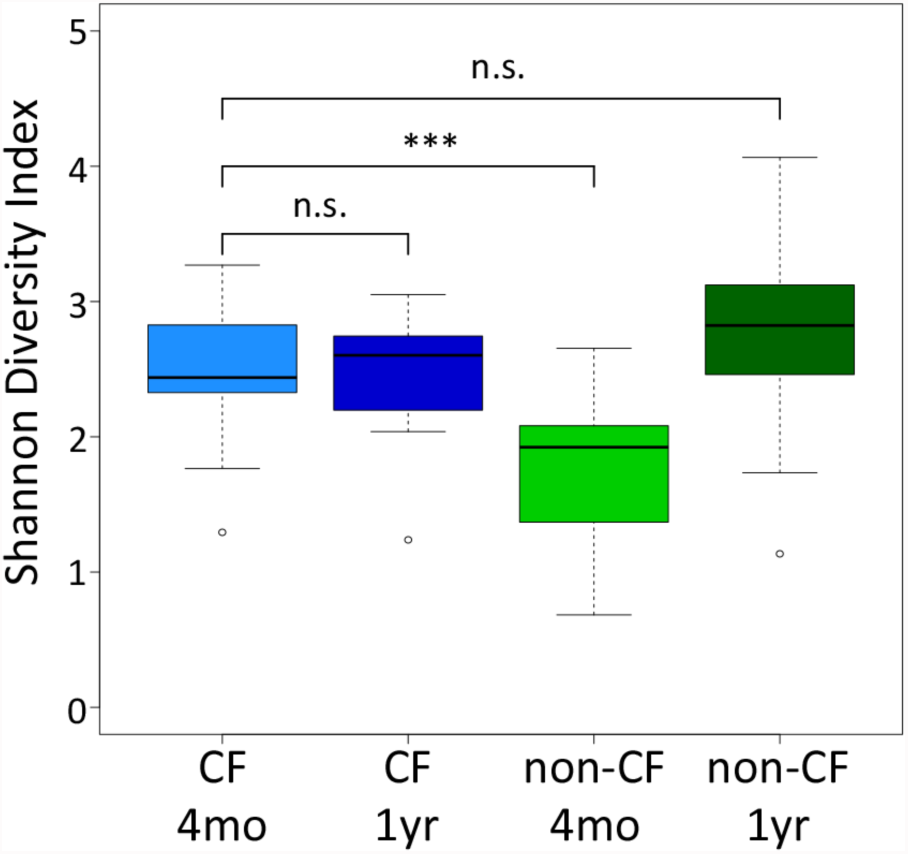
Alpha diversity varies between subjects with CF and infants without CF. Using the Shannon diversity of samples from subjects with CF at 4 months of age as the reference group in a generalized linear model, samples from CF subjects at 1 year of age are not significantly different from the same cohort at 4 months (p = 0.87). Samples collected for the comparison group (non-CF) at 4 months of age have significantly lower alpha diversity than CF subjects at the same age (p = 0.00025). While there is a trend toward increased diversity for the non-CF comparison group at 1 year of age compared to CF subjects, this trend did not reach significance (p = 0.091).

### The gut microbial communities of infants with CF differ from non-CF controls subjects over the first year of life

We next analyzed the beta diversity (microbial community composition) of the infants with CF in comparison to the non-CF comparison group. The beta diversity of patients with CF and non-CF controls at 4 months is significantly difference (Fig. 2A; PERMANOVA of generalized Unifrac distances; p = 0.002). Interestingly, despite the observation that the alpha diversity of the subjects with CF and the comparison group is not different at 1 yr, these stool samples show a significant difference in beta diversity (Fig. 2B; PERMANOVA of generalized Unifrac distances; p = 0.001). Furthermore, the similar alpha diversity of patients with CF at 4 months and non-CF children at 1 year is not reflected by a similar microbial community, that is, the beta diversity of these samples is significantly different (Fig S1; PERMANOVA of generalized Unifrac distances; p = 0.001).

**Figure 2.**
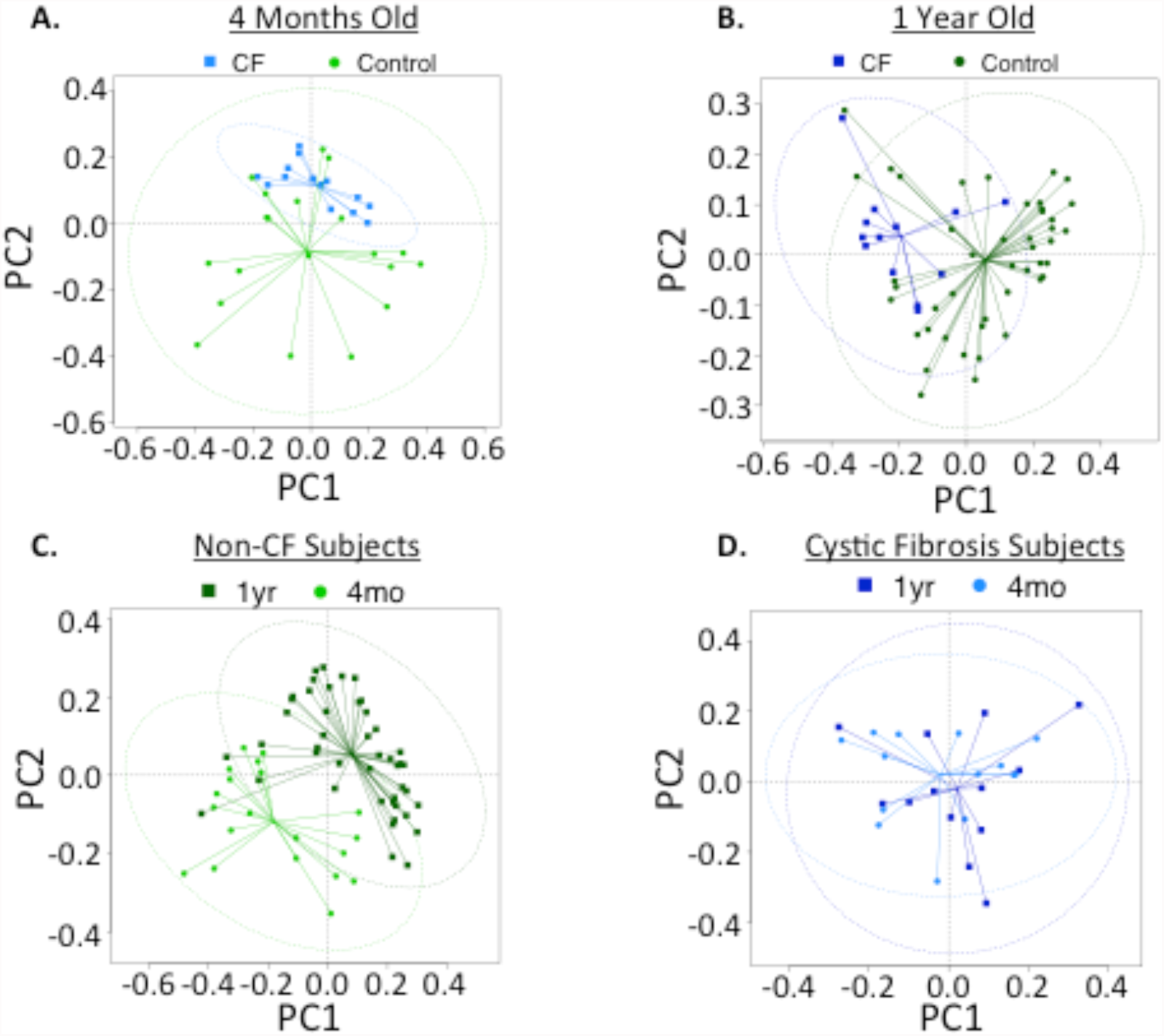
Stool microbial profile remains stable but differ between the CF and the non-CF comparison group over the first year of life. PCoA plots generated from Generalized UniFrac distances between samples. **A**. At 4 months of age samples from CF subjects cluster separately from the non-CF comparison group (PERMANOVA p = 0.002). **B**. At 1 year samples from CF subjects continue to cluster together and separately from samples provided by the non-CF comparison group (PERMANOVA p = 0.001). **C**. Samples from the non-CF comparison group at 4 month cluster separately from samples provided at 1 year (PERMANOVA p = 0.001). **D**. Samples provided by CF subjects at 4 months and 1 year do not cluster separately from each other (PERMANOVA p = 0.396).

Consistent with the significant difference in alpha diversity for non-CF children at 4 months and 1 year, there is a significant difference in beta diversity between these samples (Fig. 2C; PERMANOVA of generalized Unifrac distances; p = 0.001). For children with CF, the similar alpha diversity at 4 months and 1 year is also reflected by the clustering of communities, with no significant difference in beta diversity (Fig. 2D; PERMANOVA of generalized Unifrac distances; p = 0.396).

### Alterations in the beta diversity of stool of infants with CF are driven by modest changes in microbes for patients at 4 months of age

To further explore the basis of the differences in beta diversity between subjects with CF and non-CF controls, we investigated differences in individual OTUs, assigned at the genus level, when possible, or otherwise at the next lowest taxonomic level. Selecting the top 25 OTUs in each data set resulted in a collection of microbes that almost perfectly overlapped, and captured the top 20 OTUs for each cohort in terms of relative abundance (Fig. 3). Using a Wilcoxon rank-sum test to compare the relative abundance of genera present at 4 months, the abundance of *Bifidobacterium* is significantly higher in the non-CF controls while *Streptococcus, Veillonella*, and *Clostridium* are significantly higher in subjects with CF (Fig. 3). Three lower abundance groups (~1% relative abundance for most samples) are also significantly different: *Faecalibacterium* and two unassigned genera in the order Clostridiales. None of the other nineteen most abundant OTUs are significantly different at 4 months between these cohorts, thus only 6/25 microbes differ at the 4 month time point. By one year of life, there are more significant differences between the cohorts, with 11/25 showing a significant difference in relative abundance between these cohorts (Fig. S2). For example, *Bacteroides* and *Faecalibacterium* are significantly higher in the non-CF comparison group. There is no significant difference in *Bifidobacteria* levels between the cohorts at 1 year, despite the significant difference at 4 months (Fig. 3). Thus, particularly early in life (i.e., at 4 months), we find that the shift in microbes is primarily driven by a small subset of genera.

**Figure 3.**
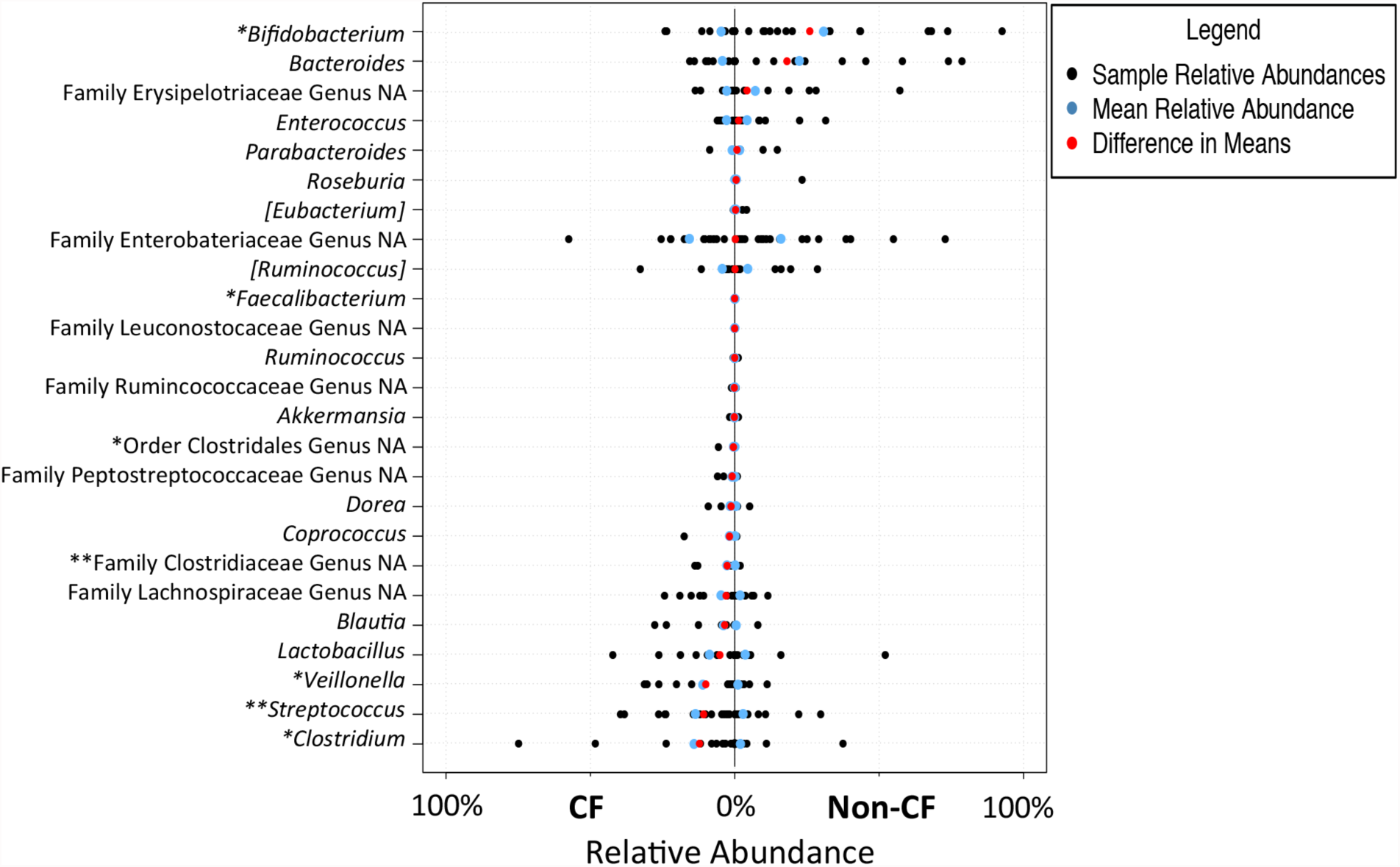
Comparison of relative abundances of genera for subjects with CF and the non-CF comparison group at 4 months. Relative abundance of the top 25 taxa present in stool samples. At 4 months there were 14 samples from CF subjects and 20 samples from the non-CF comparison group. *, **, and ***, indicate significantly different taxa by the Wilcoxon rank-sum test after Bonferroni correction at the p<0.05, p<0.01, and p<0.001 levels respectively. Genera in brackets are the taxonomic assignments suggested by the Greengenes database based on whole genome phylogeny. Abbreviations: NA, not assigned.

### Microbes associated with immune programing are lower in patients with CF compared to infants without CF

Given that subjects with CF demonstrate a lower abundance of *Bacteroides* and/or *Bifidobacterium* at 4 months and/or 1 year than the comparison group, a more indepth analysis of these two keystone immune programming genera was undertaken. While both cohorts have individual subjects with low abundance of *Bacteroides* or *Bifidobacterium*, a comparison of the relative abundance of each microbe at each sampling time revealed that for the CF cohort at both 4 months and 1 year, there were a higher percentage of patients with <1% relative abundance of these microbes and a lower maximal relative abundance compared to the non-CF comparison cohort (Figure S3).

Given the importance of *Bacteroides* and *Bifidobacterium* in immune programming (23, 24), it was unexpected that even for the non-CF comparison cohort, some individuals had <1% of these microbes. To investigate this observation further, the relative abundance of *Bacteroides* versus *Bifidobacterium* on a subject-by-subject basis for both cohorts was plotted (Fig. S4). For the non-CF cohort, many subjects with low *Bacteroides* showed a high relative abundance of *Bifidobacterium*, and visa versus (Fig. S4A). There was also a group of non-CF patients with an intermediate level of each microbe and ~50% of samples from the non-CF cohort had >40% relative abundance of *Bacteroides* and/or *Bifidobacterium*. In contrast, for the CF cohort many of the subjects clustered in a group with <20% of each microbe (Fig. S4B), and there is only 1 subject with >40% of either *Bacteroides* or *Bifidobacterium* (~5% of the samples analyzed).

### Subjects with CF and non-CF controls demonstrate differential enrichment of predicted metabolic functions

Relative to the patients with CF, the non-CF comparison infant’s microbial community was associated with increased in functions connected with β-lactam resistance, polyketide sugar unit biosynthesis, and carbohydrate metabolism (Fig. S5). In contrast, subjects with CF showed enrichment in processes that include restriction enzymes, streptomycin biosynthesis, penicillin and cephalosporin biosynthesis, and galactose metabolism (Fig. S5). Our analysis of the 4 month samples showed no significant enrichment of pathways for either cohort, despite the significant differences in abundance of several OTUs (Fig. 3, S3).

### Apical application of *Bacteroides* to intestinal epithelial cells reduces basolateral release of IL-8

A recent report examining the impact of *Bacteroides* in intestinal epithelial provides a clue to the role of *Bacteroides* in reducing systemic inflammation (25). Some *B. fragilis* strains produce enterotoxins that can stimulate the production of IL-8. Cho and colleagues showed that exposing the apical face of intestinal epithelial cells to such a toxin resulted in the increased production of IL-8 from the basolateral face of the same cell monolayer. These data indicate that impacts to the host cells in the lumen of the intestine can have systemic impacts on cytokine production. Thus, we assessed whether apical application of *Bacteroides* spp. can reduce the basolateral production of IL-8.

To investigate this mechanisms, we performed an experiment whereby Caco-2 cells were grown on transwell filters as polarized monolayers, then inoculated on the apical face with ~3.5 × 10^7^ CFU *Bacteroides thetaiotaomicron* (*B. theta*). *B. theta* was <u>not</u> introduced into the basolateral compartment. The IL-8 produced in the apical and basolateral compartments of the transwells were measured after 24 hrs of exposure to the bacteria. We found apically inoculated *B. theta* significantly reduced both the apical and basolateral production of IL**-**8 (Fig. 4).

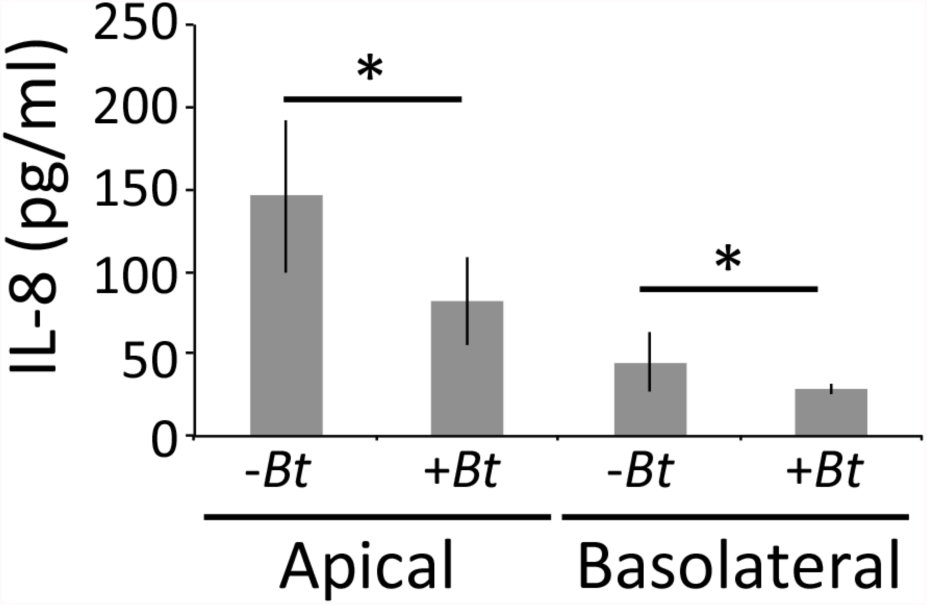
Figure 4. *B. theta* reduces IL-8 production by intestinal epithelial cells. Levels of IL-8, determined by ELISA, and indicated for Caco-2 cells grown on transwell filters for ~4 weeks. After addition of *B. theta* VPI 5482 (*Bt*) to the apical compartment of the transwell, the filters were incubated anaerobically for 3 hrs, and then the added suspension of bacteria was removed and replaced with fresh medium. Filters were returned to anaerobic incubation overnight. Twenty four hours after application of *B. theta*, supernatants were collected and IL-8 levels measured via ELISA assay (PromoKine). To determine CFU, filters were scraped and plated (n=6); at the end of the experiment there were ~2 × 10^9^ ± 4 × 10^8^ CFU *B. theta*/well associated with the apical face of the monolayer. *p < 0.05 by students t-test compared to the condition with no *B. theta* added (-*Bt*).

## Discussion

In our study, children with CF had an intestinal microbial community that was distinct from infants from the general population as early as 4 months, and despite a convergence of alpha diversity by 1 year between these cohorts, the composition of the communities remained distinct at 1 year of life. Furthermore, the altered microbiome of infants with CF did not change markedly across the first year of life. At 4 months of age, only 5/25 genera identified differed between the cohorts, but among those genera depleted at 4 months are those microbes with key immune modulatory roles. By one year of age, while the alpha diversity of the communities for the CF and the comparison cohort are not statistically significantly different, nearly half of the genera differed in relative abundance, indicating that individual taxa continue to be distinct between these two cohorts. This distinct microflora is reflected by differences in several metabolic pathways at one year.

Importantly, our previous data (15, 16) and the work here indicate a link between intestinal microbial communities and airway disease. Thus, we hypothesized that there must be a way for local effects in the intestine (i.e., altered microbiota) to impact systemic levels of inflammation and/or a gut-derived immune response, which likely contributes to airway exacerbation (26-28). Our in vitro data show that *B. theta* applied to the apical face of an intestinal cell line can reduce IL-8 secretion from both the *apical* and *basolateral* face of the cells, a finding consistent with another report in the literature that examined the impact of a *Bacteroides*-derived toxin (25). Therefore, beneficial strains of *Bacteroides* can alter systemic signals of inflammation, and thereby may impact airway disease through this mechanism.

There are several important implications of our findings. First, while enzyme replacement therapy has clearly had positive transformative impacts on CF patients (6, 29, 30), the differences we observe in the intestinal microbiota of the CF cohort indicate that these subjects still have underlying defects and suggests room for improvement in such therapeutic interventions. Additionally, our data, in combination with prior work, suggest that probiotic therapies that shift the intestinal microbiota towards a more healthful state may mitigate the pulmonary exacerbations. Several systematic reviews of randomized controlled trials of probiotics in groups with CF have shown a positive impact of probiotics on reducing frequency of CF exacerbation and improving gastrointestinal inflammation (31-33), however mechanisms behind benefits of shifting the intestinal microbiome in CF are not yet clear. Given that pulmonary exacerbations are associated with the progressive loss of lung function in this population, morbidity and ultimately mortality, reducing the frequency of such clinical episodes beginning in early life would likely assist in maintaining lung function in patients with CF.

## Materials and Methods

### Patient population

Institutional review board approval was obtained from the Center for the Protection of Human Subjects at Dartmouth College with yearly renewal of the approval. Additional detail can be found in the Supplemental Material. Eligibility criteria for the CF cohort included: diagnosis with CF prenatally or postnatally based upon newborn screening results and subsequent confirmation with sweat chloride and genetic testing. Subjects were eligible if their care for CF would be at the Dartmouth-Hitchcock Medical Center in Lebanon, NH or Manchester, NH affiliated clinics. Subjects were excluded if they had non-CF chromosomal anomalies. For the non-CF comparison group, infants were randomly selected from the New Hampshire Birth Cohort Study (NHBCS). For the NHBCS, pregnant women ages 18–45 were recruited from prenatal clinics beginning at approximately 24 to 28 weeks gestation as described: women were screened for eligibility at an initial prenatal care visit and were enrolled at 24–28 weeks gestation if they reported using water from a private, unregulated well in their home since their last menstrual period and were not planning a change in residence prior to delivery. Only singleton, live births were included in the cohort.

### Analysis of microbial populations in stool

Stool samples were analyzed as reported (16), with minor modifications, as described in the Supplemental Material.

### Statistical analyses

Differences in the microbial profiles of stool samples from 14 patients with CF and 20 comparison infants at 4 months and 13 CF patients and 45 comparison infants at 1 year were analyzed using Generalized Unifrac Distances. Distances were visualized using PCoA plots and the significance of clustering was determined through PERMANOVA. Within sample alpha-diversity was determined using the Shannon diversity index as calculated for each sample in the phyloseq R package function estimate_richness. Significant differences in the relative abundance of individual taxa were determined with a Wilcoxon rank-sum test and verified with linear models. Bonferroni correction was used to adjust for multiple testing.

### PICRUSt analysis

Open reference OTUs were removed from the OTU table using the QIIME function filter_otus_from_otu_table for use with Phylogenetic Investigation of Communities by Reconstruction of Unobserved States (PICRUSt), as reported (34). Additional detail can be found in the Supplemental Material.

### Cell line studies

Caco-2 lines were grown on transwell filters and inoculated with bacteria in the apical compartment. After 24 hours of exposure, the bacterial numbers were quantified and IL-8 measured in the apical and basolateral compartments. Additional detail can be found in the Supplemental Material.

## Acknowledgements

This work was supported by the NIH (R37 AI83256-06 and T32HL134598 to G.A.O, and P01 ES022832, P20GM104416, and, P20-GM103413) and the US EPA RD-83544201, Cystic Fibrosis Foundation (OTOOLE16G0 to G.A.O., Harry Shwachman Clinical Investigator Award, MADAN12A0, the CF-Research Deevlopment Program, STANTO07R0), and the Hearst Foundation. We thank Andy Goodman for the *B. theta* strain, and Laurie Comstock for her excellent advice and answering many questions. We thank Roxana Barnaby and the HPIC for assessing transepithelial resistance of the Caco-2 lines, and Tom Hampton from the ABBC for consulting on statistics. The HPIC and ABBC cores are supported by P30-DK117469. We also thank Devin Harbin and Michelle Tyler for their assistance.

